# Are Synaptic Clefts Directionally Oriented?

**DOI:** 10.64898/2026.01.30.702623

**Authors:** Dexuan Tang, Zhi-De Deng, Bethanny Danskin, Daniel Berger, Mark Ingersoll, Hanbing Lu, Bruce Rosen, Marom Bikson, Gregory Noetscher, Sergey Makaroff

**Affiliations:** Department of Electrical and Computer Eng., Worcester Polytechnic Inst., Worcester MA, USA; Computational Neurostimulation Research Program, Experimental Therapeutics and Pathophysiology Branch, NIMH, National Institute of Health, Bethesda MD, USA; Allen Institute for Brain Science, Seattle WA, USA; Dept. of Molecular and Cellular Biology, Harvard Univ., Boston MA, USA; University of Massachusetts Amherst, Amherst MA, USA; Neuroimaging Research Branch, NIDA, Intramural Research Program, National Institute of Health, Baltimore MD, USA; Athinoula A. Martinos Ctr. for Biomedical Imaging, Massachusetts General Hospital, Charlestown MA, USA; Department of Biomedical Engineering, The City College of New York, New York NY, USA

**Keywords:** Synaptic clefts, Synaptic orientation, Synaptic architecture, MICrONS 1 mm^3^ cortical volume, H01 1 mm^3^ cortical volume

## Abstract

Synapses are fundamental building blocks of cortical circuits, yet their geometry is typically regarded as a local property, independent of mesoscale architecture. The prevailing assumption is that synaptic clefts are isotropically oriented in space. Here, we test this assumption by analyzing approximately 117 million synaptic clefts from two independent 1 mm³ electron microscopy datasets: the human H01 middle temporal gyrus and the mouse MICrONS primary visual cortex, using three independent cleft-extraction methods. Across both volumes, we observe that synaptic cleft orientations are not randomly distributed, but instead show statistically significant and spatially coherent directional biases across cortical layers. This mesoscale anisotropy is conserved across species, yet is stronger and more consistent in human association cortex than in mouse sensory cortex, a difference that may reflect the expanded dendritic arbors and greater integrative demands of human pyramidal neurons. We propose that cleft orientation bias is a geometric consequence of the axonal and dendritic architecture that shapes synapse formation, representing a new candidate organizational feature of cortical microarchitecture with potential implications for circuit computation and neuromodulation. These findings motivate targeted physiological studies to determine whether synaptic orientation contributes causally to cortical function.

## 1. Introduction

The cerebral cortex exhibits architectural regularities that recur across species and scales, from the laminar alignment of cell bodies to the columnar organization of dendrites and axons [1],[2],[3]. These structural motifs constrain the flow of information within local circuits and shape the way cortical populations integrate activity across regions [4],[5]. Despite extensive ultrastructural characterization of synapses over several decades [6], one fundamental geometric property has remained largely unexplored: the orientation of synaptic clefts relative to cortical axes.

At the ultrastructural level, a synaptic cleft is a thin disk-shaped gap between pre- and postsynaptic membranes. To make orientation analysis tractable across large-scale electron microscopy reconstructions, we define cleft orientation along the disk’s normal axis, typically pointing from the presynaptic to the postsynaptic side. Even when clefts deviate from an ideal disk, their orientation can be quantified as the dominant averaged normal vector of the apposed membranes. The prevailing view, derived from analyses of small datasets, has been that cleft orientations are isotropically distributed at the mesoscale, a local geometric property determined by pre- and postsynaptic specializations but otherwise independent of cortical architecture [7],[8]. This isotropy assumption has never been tested at scale. The emergence of large-volume electron microscopy reconstructions now makes it possible to do so directly.

If cleft orientations are systematically biased, such anisotropy would constitute a previously unmeasured structural property of cortical microarchitecture, analogous to the preferential alignment of axons or the radial orientation of apical dendrites. Mechanistically, orientation bias is expected as a geometric consequence of the dendritic and axonal architecture that governs where and how synapses form: pyramidal neurons extend apical dendrites radially toward the pial surface, basal dendrites tangentially, and en passant axonal boutons contact these compartments at angles constrained by the local geometry. Because trans-synaptic adhesion molecules such as neurexins and neuroligins are differentially expressed across cell types, layers, and dendritic compartments [9], the distribution of synapse types and cleft orientations across cortical space need not be isotropic. Detecting such anisotropy at the mesoscale would suggest that synaptic geometry contributes to the spatial patterning of cortical circuits.

In this work, we test the isotropy assumption directly by analyzing approximately 117 million synaptic clefts from two mesoscale cortical volumes – human H01 [10],[11],[12] and mouse MICrONS [13],[14],[15],[6] – using three independent cleft-extraction methods. We observe that synaptic clefts of excitatory neurons in mammalian neocortex are not randomly distributed. Instead, they exhibit statistically significant, spatially coherent anisotropy across cortical layers. This mesoscale orientation is evident in both mouse and human cortex, consistent with an evolutionarily conserved organizational principle. The magnitude of anisotropy is significantly greater in human association cortex compared to in mouse primary visual cortex, consistent with the expanded dendritic arbors and greater integrative demands of human pyramidal neurons [16].

We also evaluate methodological artifacts arising from sample preparation and processing that can bias apparent cleft geometry and therefore require careful control. Subject to these constraints, the observed mesoscale anisotropy identifies synaptic cleft orientation as an additional, previously unmeasured axis of cortical microarchitecture. Functionally, a consistent orientation bias could shape synaptic integration across minicolumns, and thereby support lateral information flow in association areas [17],[18]. Biophysically, because synaptic clefts form nanoscale dipoles within the extracellular medium, their collective alignment could influence the generation of endogenous field potentials, as well as alter how tissue couples to exogenous stimulation [19],[20]. Together, these results can link synaptic ultrastructure to mesoscale circuit organization and to the electrical signals used to measure and perturb cortical activity.

## 2. Results

We performed a directional analysis on two mesoscale cortical volumes obtained using scanning electron microscopy (H01) and transmission electron microscopy (MICrONS), applying three independent synaptic-cleft extraction approaches and testing methods described in the *Methods* section.

- **Mesh-touch method (H01, human):** This method was applied to neuronal membrane meshes generated by Google researchers [11],[12] for excitatory neurons whose somas were located within the 1 mm³ H01 cortical volume. We processed approximately 940 pyramidal neurons from the repository (a representative subset is shown in Fig.1b,c), yielding ∼1.6 million proofread and post-processed synaptic-cleft geometries (cf. Fig.1l,m). The clefts were distributed across cortical layers as follows: L1–∼0.06 million; L2–∼0.5 million; L3–∼0.6 million; L4–∼0.2 million; L5–∼0.2 million; L6–∼0.1 million; with the remainder located in white matter.
- **Voxel-based method (H01, human):** This analysis utilized the excitatory-synapse dataset provided by the H01 team [10] in voxel format, comprising all putative excitatory synapses (∼111 million) within the H01 sample, downloaded from repository. Unlike the mesh-touch method, the voxel representation encompasses synapses originating from all neuronal processes, not solely those of pyramidal neurons.
- **Synapse-table method (MICrONS, mouse):** The publicly available MICrONS cortical column dataset [15] includes pre- and post-synaptic coordinates of all column neurons spanning all cortical layers (Figs.6,7). For each synapse, the orientation was defined by the unit normal vector pointing from the pre- to the post-synaptic side. The total number of processed clefts was ∼3.2 million (∼1.4 million in L2/L3, ∼0.6 million in L4, ∼0.4 million in L5, and ∼0.8 million in L6).

### 2.1. Synaptic clefts of H01 human brain dataset

Fig.1a shows the topology of the H01 human brain sample [10]. The *x*-axis is (approximately) directed from L6 to L1. Fig.1b is a typical postsynaptic excitatory pyramidal neuron of the sample (cyan). Red “hairs” are the associated presynaptic processes. Fig.1c is a close-up view of the apical dendrite branch located in the top right of Fig.1b. Figs.1d-k show a collection of synaptic geometries and types studied. Most typical are the axodendritic excitatory synapses of the excitatory neurons in Figs.1d-g (estimated 81% based on the mesh touch method). Among them, *en passant* boutons and synapses (Fig.1e-g) dominate terminal synapses (Fig.1d) as approximately 10:1. Figs.1h-j show less typical axosomatic connections. In Fig.1h, a part of the soma membrane is shown and axo-dendritic shaft (in Figs.1i,j) synapses of the excitatory neurons. These synapses are putatively considered inhibitory; their relative number is estimated as ∼15% again based on the mesh touch method. Fig.1k shows putative axoaxonic synapses; their relative number is estimated as approximately the remaining 4%. Those synapses are very rare and may be subject to false detection.

**Figure 1.**
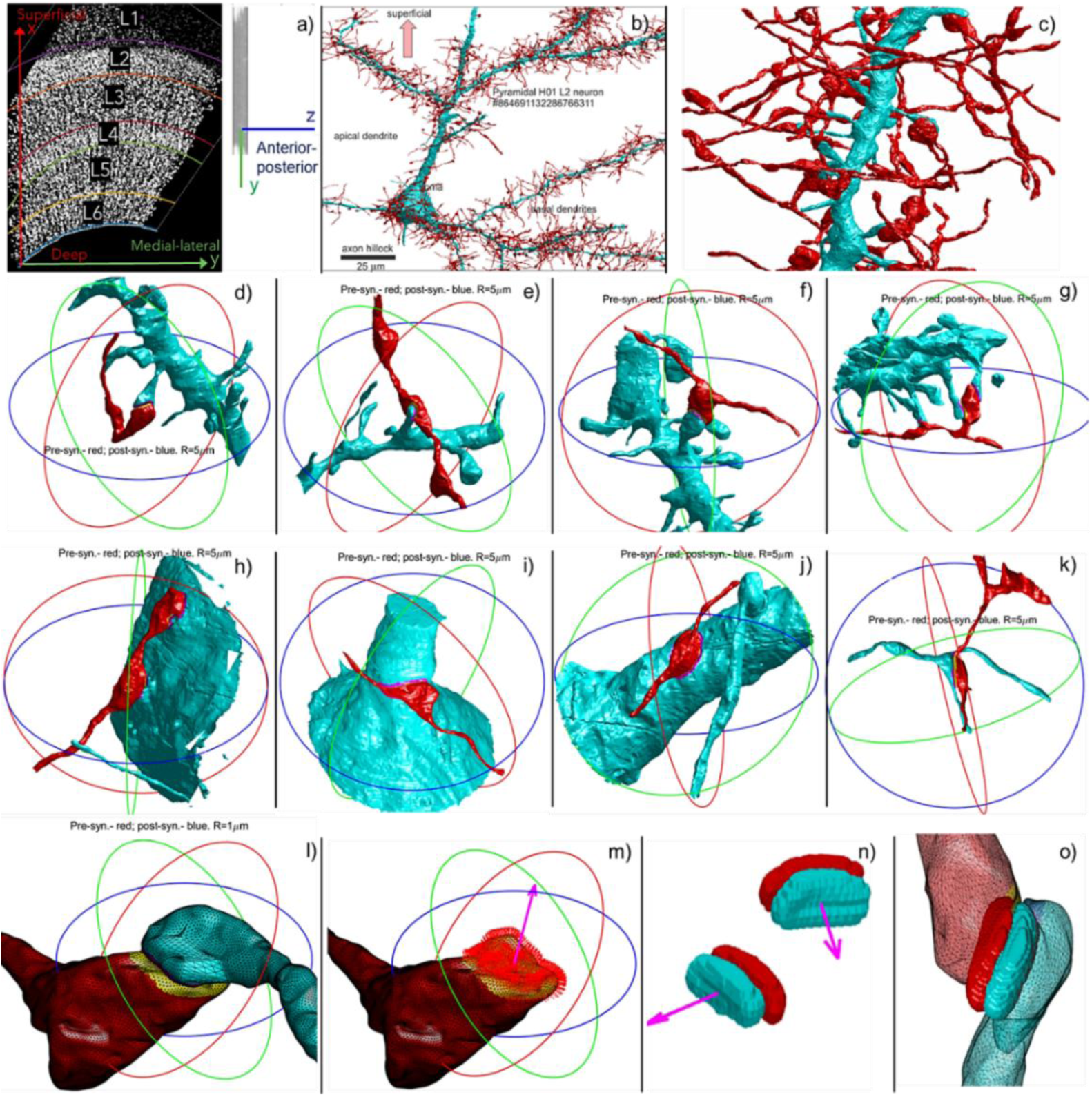
a) Topology of the H01 sample. The x-axis is directed from L6 to L1. b) Postsynaptic excitatory pyramidal neuron of the sample (blue); associated presynaptic processes (red). c) Close up view of the apical dendrite branch located top right in b). d,e,f,g) Axodendritic excitatory synapses of the excitatory neurons: en passant boutons and synapses e,f,g) and terminal synapse d). h) Axosomatic synapse with part of the soma membrane. i,j) Axo-dendritic shaft synapses. k) Rare putative axoaxonic synapses, possible false detection. l,m) Estimating synaptic cleft direction using the mesh touch method. Red normal vectors of the presynaptic membrane (yellow facets) are averaged and result in the effective directional vector (magenta). n) Estimating synaptic cleft direction using voxel data from labeling presynaptic terminals with clustered synaptic vesicles. A boundary between pre and postsynaptic voxels is extracted and subjected to Delaunay triangulation. o) Overlap of voxel and mesh touch synaptic representations (wherever possible).

### 2.2. Directional synaptic vectors of H01 human brain datasets

Using the mesh touch method (*Methods* section), normal vectors of the facets of a refined and healed presynaptic membrane (yellow in Fig.1l,m) are found first. They are shown by small red arrows in Fig.1 m. These normal vectors are further averaged (weighted by facet areas), which results in the effective directional vector of the synapse (magenta in Fig.1m). Fig.1n shows an alternative independent approach – synaptic cleft directions extracted using direct voxel data from labeling presynaptic terminals with clustered synaptic vesicles [21]. There, a boundary between pre- and postsynaptic voxels is extracted and triangulated via Delaunay triangulation, in full or partially. Normal vectors of the boundary facets are found and then averaged again weighted by facet areas. Fig.1o demonstrates overlap of voxel and mesh touch synaptic representations (wherever possible). It indicates that the voxel and mesh touch methods may generate similar, but not identical results. Additional details and direct evidence substantiating this assumption are presented in the *Discussion* and *Materials and Methods* sections.

### 2.3. “Synaptic pattern”

By analogy with the antenna radiation pattern [22], a 3D “synaptic pattern” (a 3D polar plot) has been constructed for each cortical layer of both brain samples. First, all synaptic clefts physically located within a given cortical layer are moved (only translation, no rotation) to the origin. After that, for every discrete direction in 3D characterized by a pair of polar and azimuthal angles, we introduce an elementary solid angle, *d*𝛺, enclosing this direction; the sum of all such non-overlapping *d*𝛺 is equal to 4*π*. Next, we find the total number of synaptic clefts whose primary axis or direction from Figs.1m,n – which is always oriented from pre- to postsynaptic process – lies within *d*𝛺. The synaptic pattern of the layer is constructed as a set of such absolute numbers corresponding to elementary solid angles for all different pairs of polar and azimuthal angles. The synaptic pattern is finally normalized with respect to its maximum value and is encoded via a color palette.

This definition a priori assumes that the dominant signal in the resulting synaptic pattern is coming from true orientation structure, not from a spatial sampling bias. More details and direct proof of this assumption are available in the *Discussion* section and *Materials and Methods* section.

### 2.4 Layer-by layer synaptic patterns of the 1 mm^3^ H01 human brain dataset (from association cortex) in three principal planes

Figs.2a–d, 3a–d, and 4a–d show the synaptic patterns of the H01 sample in three principal planes: *xz*-plane (Fig.2), *yz*-plane (Fig.3), and *xy*-plane (Fig.4), along with the corresponding sample topology and composition. For each plane we show in Figs.2a,3a,4a side views of the H01 sample, with cortical layers labeled from L1 (left, dark blue) to L6 (right, yellow), and representative cross-sectional planar rectangles used for plotting the Neuroglancer cell maps for each layer. We also show in Figs.2b,3b,4b the corresponding Neuroglancer slice. In all cases, all synaptic clefts located within a given cortical layer were included in constructing that layer’s synaptic pattern. We present two independent sets of patterns. The first set in (Figs.2c,3c,4c), corresponds to synaptic clefts extracted from membrane meshes using the precise mesh-touch method (Fig.1l,m and *Methods*). These patterns included both excitatory and inhibitory synapses (Fig.1d–k) of such excitatory neurons whose somas are located within the sample – approximately 1.6 million processed synaptic clefts in total. The second set in (Fig.2d,3d,4d), was obtained by processing the H01 synaptic voxel data (Fig.1n and *Methods*), which comprise 111 million voxel-based synaptic cleft geometries – all putative excitatory synapses within the sample. The same dimensionless normalized scale (0.3–1.0) is used for all human synaptic patterns.

**Figure 2.**
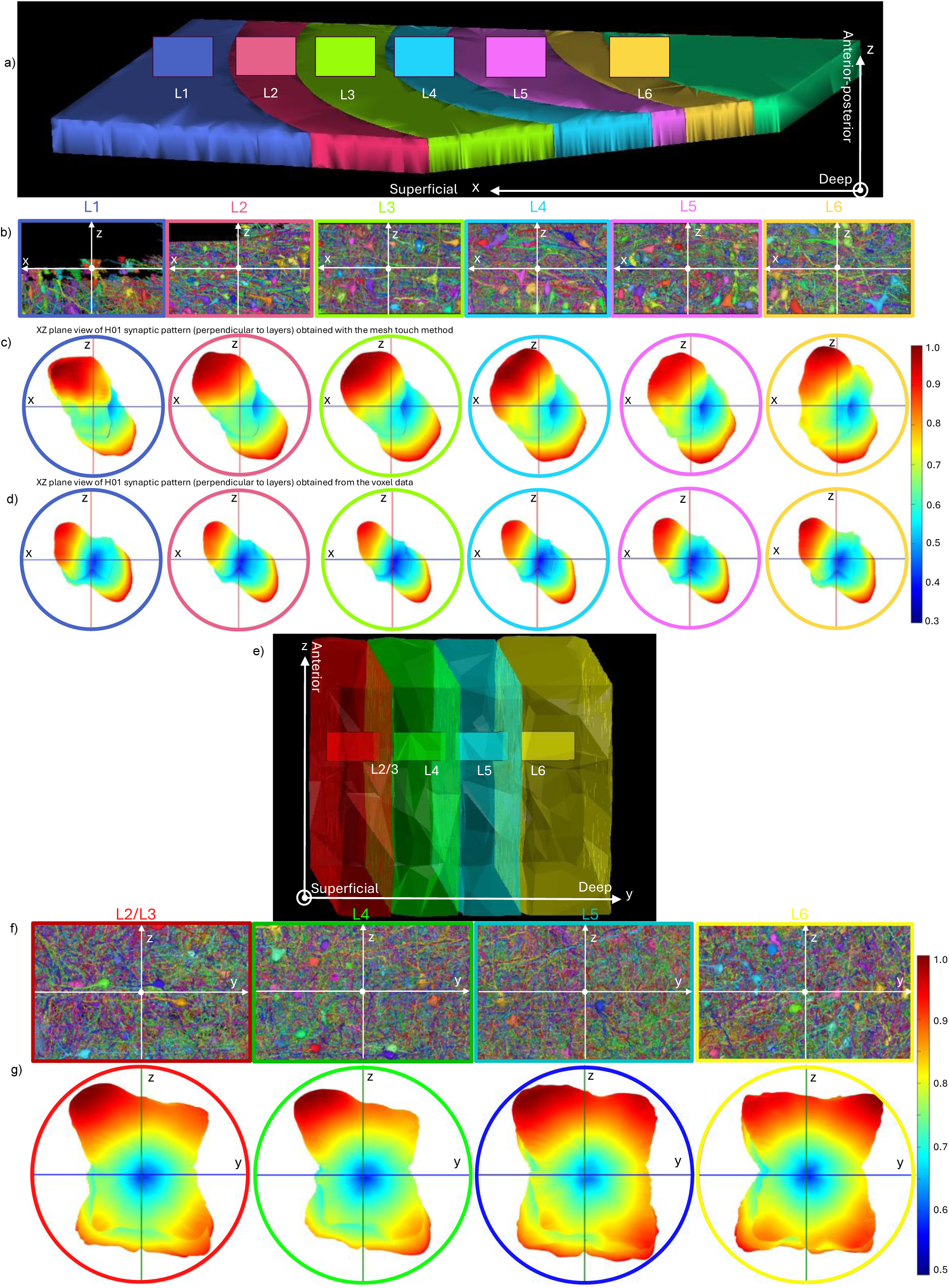
a) xz-View (side view) of the H01 sample with cortical layers labeled from L1 (left, dark blue) to L6 (right, orange) and cross-sectional planar rectangles for each layer. b) Neuroglancer slices taken along the cross-sectional planar rectangles shown in panel a). c) Layer-by-layer mesh-based synaptic patterns in the same xz-plane. d) Layer-by-layer voxel-based synaptic patterns in xz-plane. e) yz-View (side view) of the MICrONS sample with cortical layers labeled from L2/L3 (left, red) to L6 (right, yellow), and representative cross-sectional planar rectangles for each layer. f) Neuroglancer slices taken along the cross-sectional planar rectangles shown in panel e). g) Layer-by-layer synaptic patterns of the MICrONS sample in the same yz -plane. For both samples, different Cartesian coordinate systems were originally used, but the anatomical directions in the panels are the same.

**Figure 3.**
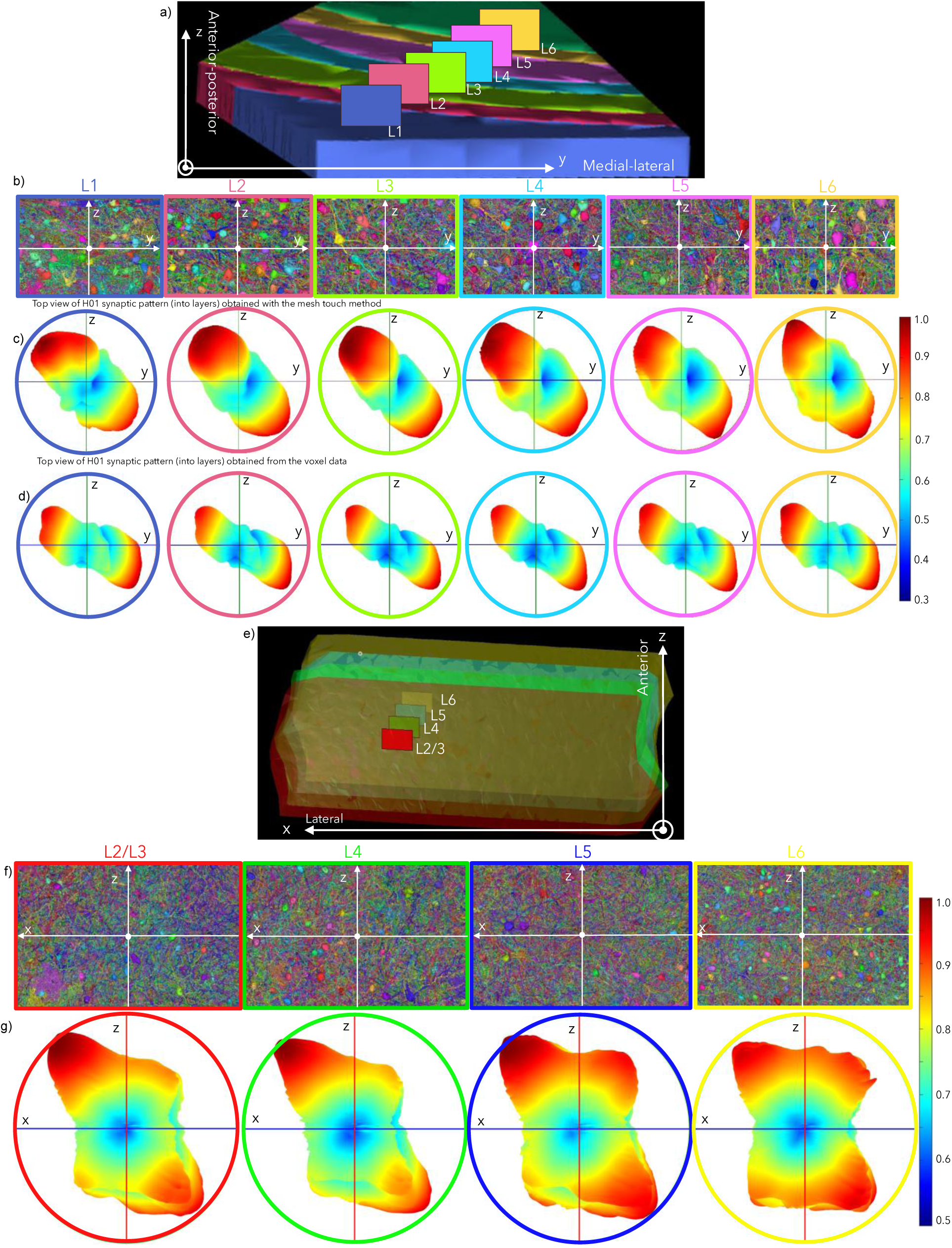
a) *yz*-View of the H01 sample with labeled cortical layers from L1 (front, dark blue) to L6 (back, orange) and cross-sectional planar rectangles for each layer. b) Neuroglancer slices in the cross-sectional planar rectangles from panel a). c) Layer-by-layer mesh-based synaptic patterns in the same *yz*-plane. d) Layer-by-layer voxel-based synaptic patterns in the yz-plane. e) *xz* -View of the MICrONS sample with labeled cortical layers from L2/L3 (left, red) to L6 (right, yellow) and cross-sectional planar rectangles. f) Neuroglancer slices in the cross-sectional planar rectangles from panel e). g) Layer-by-layer synaptic patterns of the MICrONS sample in the same *xz*-plane

**Figure 4.**
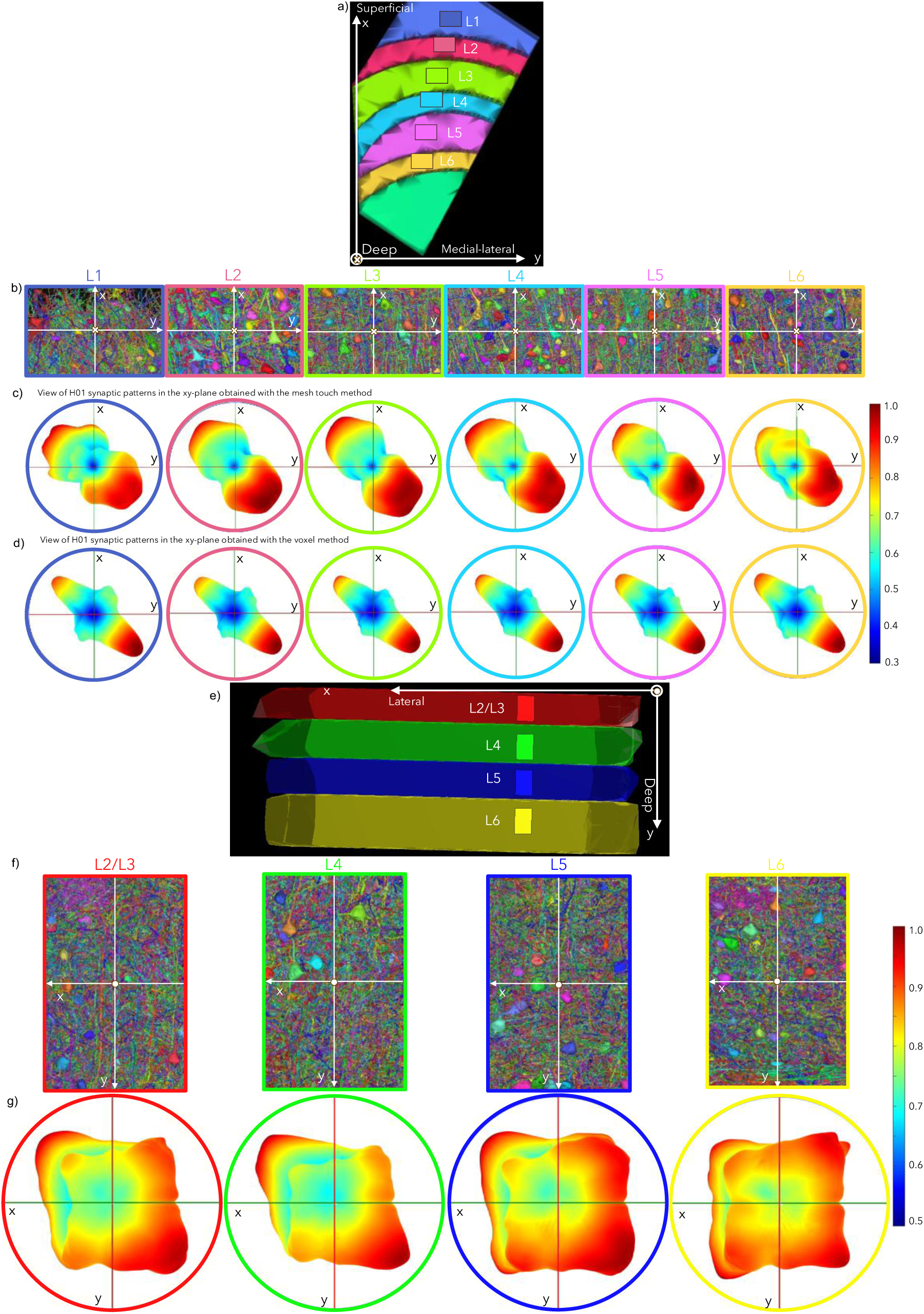
a) ***xy*** -View of the H01 sample with labeled cortical layers from L1 (top, dark blue) to L6 (bottom, orange) and cross-sectional planar rectangles for each layer. b) Neuroglancer slices in the cross-sectional planar rectangles from panel a). c) Layer-by-layer mesh-based synaptic patterns in the same ***xy***-plane. d) Layer-by-layer voxel-based synaptic patterns in the same ***xy***-plane. e) ***xy***-View of the MICrONS sample with labeled cortical layers from L2/L3 (left, red) to L6 (right, yellow) and cross-sectional planar rectangles where the Neuroglancer cell maps will be plotted) for each layer. f) Neuroglancer slices in the cross-sectional planar rectangles from panel e). g) Layer-by-layer synaptic patterns of the MICrONS sample in the same ***xy***-plane.

### 2.5 Layer-by layer synaptic patterns of the 1 mm^3^ MICrONS mouse cortical column (from primary visual cortex) in three principal planes

The same three-plane layout is shown for the MICrONS column in Figs.2e–g, 3 e–g, and 4e–g, in the order of *yz*-plane, *xz*-plane, and *xy*-plane. For each plane we show in Fig.2e,3e,4e side views of the MICrONS sample, labeled from L2/3 (left, red) to L6 (right, yellow). Representative cross-sectional planar rectangles, used for plotting the Neuroglancer cell maps, are also indicated for each layer. We also show in Figs.2f,3f,4f the corresponding Neuroglancer slice. The synaptic cleft orientations (Figs.2g,3g,4g) were acquired directly from the MICrONS database as described in the *Methods* section. The mouse brain patterns include only excitatory-neuron synapses and are based on approximately 3.2 million processed synaptic clefts of the MICrONS cortical column.

Although different Cartesian coordinate systems were originally used for human and mouse samples, we maintained consistent anatomical orientations across all three figures. Again, all synaptic clefts residing within a cortical layer were included in the construction of that layer’s synaptic pattern. The dimensionless normalized scale (0.5-1.0) is used for mouse synaptic patterns in Figs.2,3,4. It differs from that used for the human data (0.3-1.0), providing finer contrast and allowing smaller variations in the synaptic pattern to be resolved.

## 3. Discussion

### 3.1 Findings

Across all three cleft-extraction approaches, and across both species, synaptic clefts in mammalian cortex were not randomly oriented. Instead, they exhibited a conserved, statistically significant anisotropy (Figs.2–4). This is confirmed by randomization controls showing that the result is not an artifact of finite sampling (Fig.7d). The principal orientation axis did not consistently align with cortical columns but instead appeared to reflect location-specific cortical anisotropy, as visible in Figs. 2b, 3b, 4b. Orientation patterns showed moderate layer-wise variability and were bidirectional, consistent with the geometry of symmetric pre- and postsynaptic membranes. Anisotropy was present in both species, but its magnitude and layer-wise consistency were substantially greater in human association cortex than in mouse primary visual cortex. The one exception was mouse L6, where artifactual distortions associated with column curvature and increased myelination complicate interpretation.

### 3.2 Methodological robustness and limitations

The voxel-derived patterns of the H01 dataset (Figs.2d,3d,4d) exhibited the weakest variability across cortical layers and a consistent strong minimum along the anterior–posterior (*z*) direction. This minimum is not biological in origin but reflects a major methodological artifact. As explained in Fig.8, the synapse classifiers used to construct the H01 dataset were trained on EM images with high *xy*-resolution (4 nm) but substantially lower z-resolution (33 nm). Postsynaptic densities oriented within *xy*-planes were therefore detected more readily than those aligned along *z*, and this anisotropic detectability was inherited by the machine-learning classifiers. In contrast, the mesh-touch method is less affected by this bias (Figs.2c,3c,4c). A separate, sample-compression correction described in the *Materials and Methods* section was also applied.

In the MICrONS cortical column, which exhibits comparable imaging anisotropy, the synaptic orientation patterns display analogous minima (Figs.2g,3g,4g). These patterns consist primarily of a directional component superimposed on an artificial rectangular “dumbbell” feature that is most pronounced in Layer 6. Imperfections in the manually trained synapse classifiers may contribute to this artifact. Bending of the cortical column in L6 (Fig.6f) likely amplifies this distortion, while additional segmentation challenges associated with the increased myelination toward L6 may further accentuate it.

In both datasets, irregular sample geometries can potentially distort unweighted directional patterns. To mitigate this, we applied a weighted directional analysis analogous to those used in materials science and crystallography [23],[24] and, implicitly, in neuroscience [25], where local density normalization is employed to reduce anatomical bias in orientation fields. The *Methods* section details this methodology and provides a side-by-side comparison for the MICrONS dataset, yielding results that closely match those shown in Figs.2–4. A similar validation was performed for the H01 dataset, confirming that spatial sampling density does not vary systematically with direction – that is, sampling is sufficiently isotropic.

Robustness was further tested by extracting multiple spherical subvolumes from different cortical layers in both datasets and repeated the analyses. The resulting patterns were nearly identical, indicating that the dominant features in our directional histograms – except for the z-direction artifact in the H01 dataset – reflect genuine synaptic-cleft orientation structure rather than geometry-specific sampling bias. Unavoidable residual effects may also include annotation errors, meshing inaccuracies, and tissue deformations introduced during sample preparation that could not be fully accounted for. Notably, across both datasets and all three extraction methods, the reconstructed synaptic patterns converge toward a stable form even when relatively few clefts (∼25,000–50,000) are analyzed. As the number of included clefts increases, the overall anisotropic patterns remain robust and change only modestly.

A further interpretive limitation is that the two datasets originate from different cortical regions: human association cortex (middle temporal gyrus) and mouse primary visual cortex (V1). The greater anisotropy in the human sample could therefore reflect regional specialization rather than a species difference per se. Distinguishing these possibilities will require datasets from matched cortical regions across species, which are not currently available at comparable scale and resolution.

### 3.3 Possible mechanistic basis for cleft orientation

Cortical pyramidal neurons extend apical dendrites radially toward the pial surface and basal dendrites tangentially within their home layer. En passant axonal boutons, which dominate over terminal synapses in both datasets, contact these dendritic compartments at angles that are constrained by local geometry. Because the normal to the synaptic cleft is determined by the contact angle between the axonal process and its dendritic target, the distribution of cleft orientations across a cortical volume will directly reflect the distribution of dendritic orientations, which is far from isotropic.

At the molecular level, this geometric argument is supported by the known cell-type and compartment specificity of trans-synaptic adhesion molecules. Neurexin splice isoforms are differentially expressed across neuron types and brain regions in patterns that are temporally and spatially regulated [9],[26]. Neuroligin isoforms are similarly segregated: Nlgn1 localizes predominantly to excitatory synapses, Nlgn2 to inhibitory synapses, and Nlgn3 to subsets of both [27]. Because different neuroligin isoforms are concentrated on different dendritic compartments, and because these compartments have preferred orientations relative to cortical axes, the aggregate distribution of cleft orientations will inherit the geometric bias of the dendritic arbor. No field-sensitivity or orientation-specific molecular signaling is required; the anisotropy emerges from the spatial distribution of where synapses form.

One supporting line of evidence is recent work on the mouse brain synaptome [28] which demonstrated that synapses are not uniform but exhibit region- and layer-specific patterns of molecular composition and morphology. Zhu and colleagues [28] showed that these spatial gradients in synapse diversity shape network topology and functional connectivity, providing evidence that synaptic properties contribute to mesoscale circuit organization. Our study extends this principle by identifying orientation as an additional dimension of synaptic architecture. Whereas Zhu et al. highlighted diversity in molecular identity and spine morphology, our results indicate that the geometry of the synaptic cleft itself is patterned relative to cortical axes. Together, these findings suggest that synaptic microstructure may contribute to cortical computation along multiple orthogonal dimensions: molecular diversity, morphological specialization, and spatial orientation. This framework positions synaptic architecture as a multiscale substrate for evolutionary and functional differences in cortical processing.

The anisotropy was present in both mouse and human cortex, establishing it as a conserved rule across species. However, the magnitude and layer consistency of the bias were greater in human than in mouse cortex. This difference aligns with broader comparative neuroanatomy: human cortex is thicker, and pyramidal neurons possess more extensive dendritic arbors, with apical and basal branches orders of magnitude longer than in mouse, while human neurons also receive substantially more synaptic inputs per dendritic segment [29],[30],[31].

We hypothesize that stronger anisotropy in human association cortex may reflect (i) evolutionary scaling of synaptic geometry to support greater integration across cortical columns and/or; (ii) specialization of the brain region in which it occurs – as pointed out, for example, in Barbas et al 2022 [32].

Recent cross-species imaging work reinforces the first interpretation. Liu et al. demonstrated that cortical architecture exhibits species- and layer-specific trajectories [33]: in both humans and mice, deep layers thin with age while layer IV becomes relatively thicker and more myelinated, but the magnitude and functional implications of these changes differ between species. Their findings underscore that laminar microstructure is not fixed but evolves in ways that support distinct modes of integration. In parallel, our observation that synaptic clefts are more strongly and consistently anisotropic across layers in human cortex suggests that orientation bias is another measurable property along which cortical microarchitecture scales with integrative demands. Taken together, these results argue that both laminar architecture and synaptic orientation anisotropy represent species-sensitive structural adaptations that enhance cross-columnar and cortico–cortical communication.

On the other hand, there could be significant differences in the columnar and laminar organizations between different brain regions. Barbas et al. noted that feedforward connections, where the signal travels from earlier to later processing cortices, have a vertical organization. This is in contrast with feedback connections, where the signal travels from later to earlier processing areas have a more laminar organization. This could also be a possible explanation for the difference in anisotropy observed in the human and mouse data, as the two regions (association cortex vs primary sensory cortex) may have a preference in the direction which the signal travels [32].

Several factors may contribute to the observed anisotropy. At the cellular level, the expanded dendritic arbors of human pyramidal neurons increase the likelihood of tangentially aligned synaptic contacts. Developmental guidance signals and structural scaffolds, including radial glia and molecular gradients [34],[35], may impose constraints that orient presynaptic and postsynaptic membranes relative to cortical axes. At the tissue level, interactions with extracellular matrix could reinforce preferred orientations. Together, these cellular and developmental mechanisms may embed orientation bias as a byproduct of cortical architecture.

Orientation bias may enhance lateral signal integration across minicolumns and modules and vice versa. In association cortex, such integration is essential for combining multimodal inputs and supporting higher-order cognition. By aligning synapses preferentially in tangential planes, cortical microcircuits could potentially facilitate cross-columnar communication while reducing redundancy along the radial axis. This aligns with evidence that human cortex favors cortico-cortical connectivity emerging from supragranular layers [17],[18]. At a broader scale, synaptic anisotropy could shape the propagation of activity across cortical networks, influencing oscillatory dynamics and inter-areal coordination.

Bodor et al. (2025) [36], analyzing the same MICrONS dataset, showed that layer 5 thick-tufted pyramidal neurons (extratelencephalic cells) in mouse visual cortex predominantly form local synapses onto inhibitory interneurons, which reciprocally inhibit them, while their longer-range projections target excitatory populations such as L6 pyramidal neurons and L5-IT cells. This architecture implies local competition and long-range coordination, consistent with predictive coding and the generation of gamma oscillations. Notably, Bodor et al. highlight that such local inhibitory targeting differs from cat and monkey cortex, where excitatory neurons more commonly synapse onto other local excitatory cells. These findings are in line with our observation that synaptic cleft orientation anisotropy is weaker and less consistent in mouse cortex than in human cortex: both results point to a circuit-level strategy in rodents that emphasizes inhibitory balance locally, in contrast to the more integrative excitatory connectivity characteristic of higher order animals.

In conclusion, our analyses suggest that synaptic clefts in mammalian cortex are not randomly oriented but may exhibit a conserved anisotropy at the mesoscale. This orientation bias was evident in both human and mouse datasets, yet it appeared stronger and more consistent in human association cortex. We propose that anisotropy in cleft orientation may reflect the geometry of axonal and dendritic arbors and the developmental scaffolds that shape them, and that its amplification in humans could relate to the expanded integrative capacity of association cortex. Furthermore, the collective alignment of synapses may influence how cortical tissue interacts with endogenous field potentials and with externally applied brain stimulation.

At the same time, we emphasize that the present findings must be interpreted with caution. Both datasets are subject to methodological artifacts, resolution anisotropies, and potential biases in segmentation and classification. The observed anisotropy may therefore, at least in part, reflect technical or sampling effects rather than an intrinsic biological property. Definitive validation will require independent datasets acquired at truly isotropic voxel resolution (∼10×10×10 nm), improved AI-based reconstructions with minimized geometric distortion, and targeted physiological and computational studies to link structure with function.

Despite these limitations, the possibility that a conserved, orientation-dependent organization exists across mammalian synapses is intriguing. By revealing a candidate mesoscale feature of synaptic architecture and by framing it explicitly within its methodological constraints this work aims to motivate deeper experimental and theoretical investigations into how microstructural geometry contributes to cortical computation and neuromodulation.

## 4. Materials and Methods

### 4.1 Synaptic cleft extraction and processing for H01 dataset

Chemical synaptic clefts were extracted from a micrometer thick slab of human cortex (H01) from the anterior part of the middle temporal gyrus, with a total volume of just over one cubic millimeter. The sample was removed to gain access to an epileptic focus in the underlying hippocampus [10]. In its longest dimension, the sample spans all cortical layers (6 through 1) in the *x*-direction (from deep to superficial) but is relatively narrow (∼175 µm) in the anterior-posterior (*z*) direction. By light microscopy-based neuropathological examination, the sample was deemed normal [10]. To align the cortical direction with the *x*-axis, the entire sample was rotated by 35° about the *z* -axis, resulting in the orientation shown in Fig.1a of the main text.

#### Mesh touch method for extracting cleft directions

The starting point of our analysis is the synaptic repository [11] of the H01 sample publicly available via Google Cloud storage [12]. Two parts of the repository were used: (i) surface membrane meshes created by a Google meshing team in the STL format for individual pyramidal excitatory postsynaptic neurons with adjacent presynaptic neuronal (predominantly axonal) processes (as shown in Fig.1b,c) and; (ii) the corresponding CSV synaptic tables (one such table per postsynaptic neuron) that list pre- and postsynaptic surface mesh IDs, as well as the coordinates for each side (side center) of the synaptic cleft. In this analysis, we did not differentiate between excitatory and inhibitory synapses for the membrane surface meshes although the dominant axodendritic synapses are mostly excitatory.

Based on the presynaptic and postsynaptic center positions, and using manipulations with the available triangular surface meshes, we have extracted the membrane topology near the cleft within approximately 1 µm from its center as shown in Fig.1l). We have further extracted both pre- (yellow in Fig.1l) and postsynaptic (magenta in Fig.1l) membrane interfaces. Those are defined here as separated from each other by at most 100 nm. A test was done when reducing this value to 75 nm and very similar results were obtained. The corresponding meshes were further refined by 1:4 manifold triangular subdivision followed by surface-preserving Taubin smoothing [37].

In this way, we processed 944 pyramidal neurons of the repository which resulted in a total of 1,566,358 proofread and postprocessed synaptic clefts. The synaptic clefts themselves are mostly located between layers L2 to L6 (L1: 56,649; L2: 482,282; L3: 559,968; L4: 165,621; L5: 172,241; L6: 99,995; the rest is in white matter).

As shown in Fig.1m, for every yellow presynaptic membrane interface, we found the total average outer normal vector by averaging the normal vectors of the respective triangular facets (weighted by facet areas) of the presynaptic interface. This average normal vector (magenta arrow in Fig.1m) has been chosen as the (pre) synaptic orientation of the cleft or its primary directional axis. This axis denotes the approximate direction in which the neurotransmitters travel from pre- to postsynaptic processes.

#### Voxel method for extracting cleft directions

One potential limitation of the previous method is that it includes only neurons whose somas are located within the sample volume, thereby excluding orphaned fragments of the neuropil contained inside the dataset. To address this issue, we additionally analyzed the excitatory synaptic data provided in voxel format, which encompass all putative excitatory synapses (111,272,315 in total) within the sample. The same Google Cloud repository also contains the voxel representation of synaptic clefts spanning the entire sample volume [12]. Unlike the mesh-based approach, the voxel representation includes synapses originating from all neuronal processes, not only from pyramidal neurons.

For our analysis, we used two available volumetric renderings decompressed from the *Neuroglancer* format [38]: (i) *Volumetric E/I Synapses* that contains a voxel representation of the entire synapse (both pre- and postsynaptic sites, but the voxels are not discriminated between the two types, shown in red in Fig.1n) and; (ii) *Incoming excitatory synapses* that contains voxel representation of the excitatory postsynaptic density or PSD (shown in blue in Fig.1n). Both volumetric renderings have identical dimensions (515,892×356,400×5,249 voxels) and resolutions (8 nm per voxel in the *x* and y directions and 33 nm per voxel in the *z*-direction).

The entire volume was originally divided into ∼6.4 million regions of interests, each with a size of 1,000×1,000×150 voxels. For each region, voxels belonging to the individual synapses were separately extracted using the *ind2sub* and *bwconncomp*-based clustering method in MATLAB. After that, we analyzed every individual synaptic assembly separately. Namely, for every presynaptic voxel, we found all postsynaptic voxels within a 40 nm distance. In total, 111,272,315 excitatory synapses (all synapses available) were processed.

To ensure that no synaptic neighborhood was artificially truncated at the borders of our 1,000×1,000×150-voxel blocks, each chunk was extended by a volumetric “margin” whose thickness in X, Y, and Z matched the maximum synapse-to-synapse search radius (40 nm converted to voxels). In practice, for every region we first loaded an additional pad of voxels on all six faces, built connected-component labels using *ind2sub* and *bwconncomp*-based clustering and distance searches over the full padded region, and only after computing each synapse’s orientation did we discard any result whose centroid lay within that padding. This approach guaranteed that every potential partner voxel within one search radius (40 nm) of a presynaptic site was present in the analysis – even at chunk boundaries – while at the same time preventing any synapse from being counted more than once.

By filtering out synapses whose centroids fell in the padded margins, we retained exactly one fully correct orientation vector per synapse and avoided artifacts that would otherwise arise when a search sphere is clipped by an artificial cube face. The result is a seamless tiling of orientation vectors across the entire volume, with no edge-effects and no overlap between adjacent blocks. Crucially, although we discard every synapse whose centroid lies in the padded “margin for each processed region,” no synapse is ever lost. The reasons being:

i. Every synapse appears fully inside at least one padded block. By extending each 1,000×1,000 × 150-voxel chunk by a “margin” equal to the 40 nm search radius on all six faces, even a synapse whose true centroid lies exactly on the boundary of one central block will lie well inside the padded volume of its neighbor. In that neighboring block the synapse never touches the padded-minimum faces and so is processed with its full 40 nm neighborhood intact.
ii. We record only synapses whose centroids lie in the original (unpadded) block.

After computing every orientation vector in the padded region, we tested each synapse’s global centroid against the global coordinate ranges of each region and saved only those that fall inside. Since a given centroid can lie in exactly one block’s central region, every synapse is committed exactly once – and only once.

#### Delaunay triangulation-based method for extracting cleft directions from voxel data

Our method is illustrated in Fig.5a–c. Extraction of synaptic cleft orientation from the original voxelized synaptic data (Fig.5a) begins by identifying the boundary between the two voxel sets shown in Fig.5b. This boundary is then triangulated (Fig.5c), after which the normal vectors of the resulting triangular facets are computed and subsequently averaged to determine the cleft direction. In cases where Delaunay triangulation performs poorly due to complex synapse geometries, partial triangulation is applied to multiple subsets of the synaptic boundary. This procedure was used to extract the orientation of all synaptic clefts analyzed in this study.

**Figure 5.**
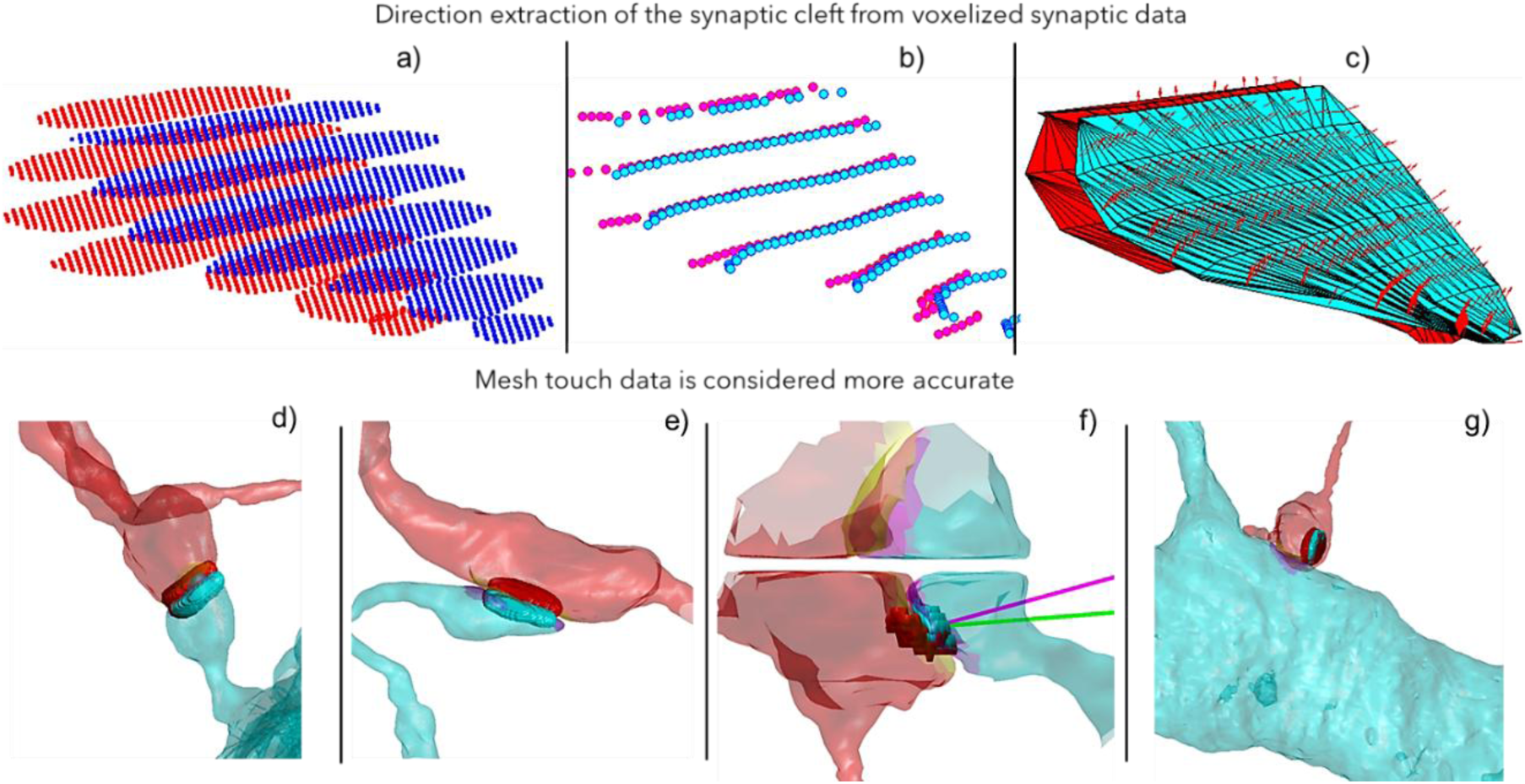
a,b,c) Illustration of Delaunay-triangulation method used for extracting cleft directions from original voxel data in a). In b), a boundary between two voxel sets is extracted, which is further triangulated in c). Normal vectors of the triangular facets are then found and finally averaged. d,e,f,g) Correlations between voxel and mesh touch synapse representations. Frequently, both methods agree reasonably well as in d,e). However, there are many cases where the voxel-based method is missing a part of the synapse as in f). As a result, the cleft direction shown by the respective arrow becomes different. In certain cases, the cleft is identified by the voxel method totally incorrectly as in g).

#### The mesh-touch method may be more accurate than the voxel-based method

In most cases, the two methods produce reasonably consistent results, as illustrated in Fig.5d,e. However, there are instances in which the voxel-based method fails to capture portions of a synapse, as shown in Fig.5f. Consequently, the estimated cleft direction (indicated by magenta and green arrows in Fig.5f) deviates between the two methods. In other cases, the synaptic cleft is only partially identified or misclassified altogether (Fig.5g). The exact frequency of such occurrences remains undetermined.

### 4.2 Sample compression correction for the H01 dataset

It is important to consider that artifacts introduced during H01 sample preparation may influence the measured synaptic orientations. The geometry of the H01 dataset originates from multibeam scanning electron microscopy (ATUM–mSEM) rather than transmission electron microscopy (TEM). In the ATUM workflow, each 30–33 nm section is cut, mounted on Kapton tape, and scanned using 61 electron beams at 4 × 4 nm pixel resolution over an area of approximately 4.5 mm². Mechanical pickup during this process anisotropically flattens each section: the authors of [10] reported nearly 28% compression along the axis perpendicular to the knife edge (*x*-axis) and only ∼0.3 % along the axis parallel to it (y-axis) (see Supplementary Information of Shapson-Coe *et al.*, 2024 [10]).

By contrast, the serial-section TEM used for the MICrONS dataset images entire sections in transmission. In this technique, dimensional distortions are typically isotropic—on the order of 2–5 % shrinkage—arising from resin polymerization and drying, rather than the directional flattening inherent to tape-collect SEM [39],[40].

To correct for this effect (not accounted for in the publicly available datasets), we expanded the H01 sample by 28% along the *x*-axis and correspondingly reduced the *x*-components of all synaptic normal vectors by the same proportion.

### 4.3 Synaptic cleft extraction and processing for MICrONS dataset

#### Using synaptic tables

We analyzed the *Minnie65* subset of the MICrONS dataset [6], representing the primary visual cortex of a mouse. This subset contains a cortical column comprising 1,356 neurons and measuring approximately 200 µm in diameter. The column spans all cortical layers and is shown in Fig.6 (axons – cyan; apical dendrites – magenta; basal dendrites – blue; somas – orange). It occupies a substantial portion of the overall sample volume and exhibits a slight curvature in layer 6. The *Minnie65* dataset was accessed via CAVEclient (Python 3.11.9), using materialization v1181, which was the most recent public release at the time of analysis. For the present study, we selected all excitatory neurons (1,097 in total) within this cortical column.

**Figure 6.**
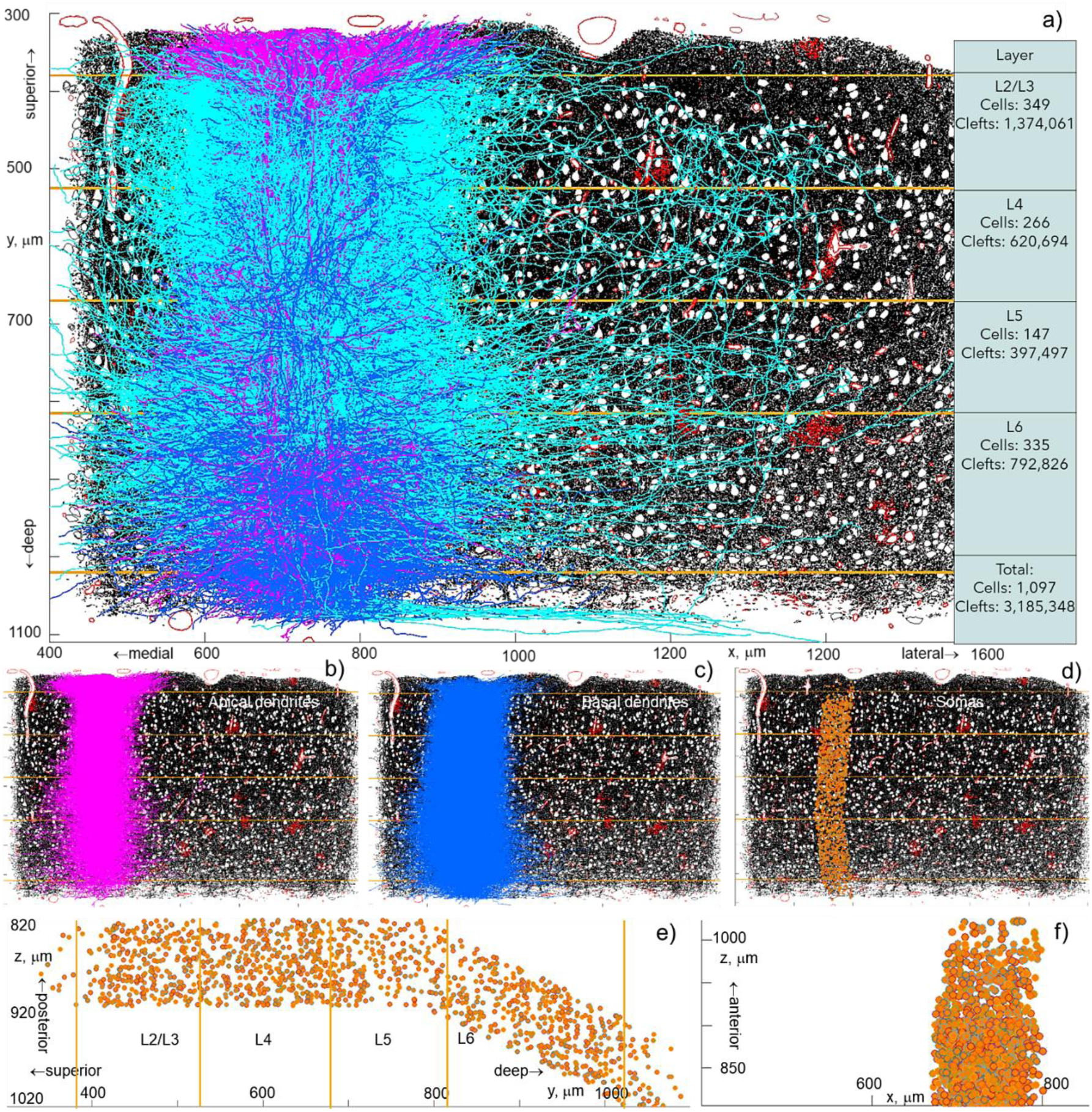
a) Side view of the IARPA Phase III MICrONS V1 cortical column showing 1,097 excitatory neurons overlaid on a cross-section of the *Minnie65* cortical volume at *z* = 937 *μm*. Cell membranes are shown in black, the vasculature in red, and somas in white. Approximate layer boundaries are indicated by orange lines. Axons are shown in cyan, apical dendrites in magenta, and basal dendrites in blue. The table on the left quantifies the number of cells with somas located within each layer and the number of synaptic clefts physically contained in that layer. b,c,d) Separate visualizations of apical dendrites (b), basal dendrites (c), and somas (d). e,f) Illustration of cortical column curvature in the *yz*-plane, with somas shown in orange.

The synapse table (synapses_pni_2) includes pre- and post-synaptic side point positions. This information was used for determining effective synaptic cleft orientations (and thicknesses if necessary). Additionally, the manual cell types (V1 column) table (allen_v1_column_types_slanted_ref) was referred to for determining detailed cell type information necessary for filtering and categorizing downloaded cell data. The root IDs were extracted and organized by layer into the four categories (L2/3, L4, L5, and L6). These extracted root IDs were used to filter the synapse table query by specifying them as post_ids. Cell data from the synapse query were downloaded as csv files and organized by layer. The total number of processed clefts is ∼3.2 million (1.4 million in L2/3, 0.6 million in L4, 0.4 million in L5, and 0.8 million in L6). To determine the orientation of each cleft, unit normal vectors were constructed pointing from the pre- to the post-synaptic side. No differentiation between excitatory and inhibitory synapses was made.

#### Layer-by-layer separation of synaptic clefts

Approximate layer boundaries (Fig.6a) were determined based on local nuclear features [40]. In this region of the visual cortex, layers 2 and 3 are not well demarcated and are therefore collectively referred to as Layers 2/3. The generated meshes delineate the borders between Layer 1, Layers 2/3, Layer 4, Layer 5, Layer 6, and the white matter. Manually drawing layer boundaries in approximately the correct anatomical locations was considered sufficient for the purposes of this analysis. All synapses were then assigned to their respective layers according to spatial position, and three-dimensional synaptic orientation patterns were subsequently constructed for each layer as described in the Results section.

### 4.4 Weighted versus unweighted synaptic patterns

To minimize the effect of the sample shape and other inhomogeneities, we applied the weighted directional analysis used in materials science and crystallography [23],[24] and, implicitly, in neuroscience [25]) where the authors apply local density normalization in histological mapping to avoid anatomical bias in orientation fields. In essence, the previous definition of the synaptic pattern was modified by

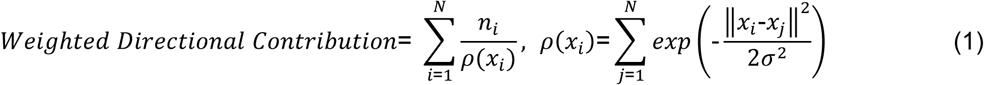

Here, *x*_*i*_ are positions of *N* synapses, *n*_*i*_ are the cleft orientation vectors, *ρ*(*x*_*i*_) is the local spatial density of the synaptic clefts which is found as a probability density estimate. This method in particular corrects for a non-uniform sample geometry.

The results presented in Figs. 2, 3, and 4 of the main text were re-computed using the new definition (Eq. 1) with a moderate standard deviation of 0.3, and the resulting patterns were compared across datasets. The outcomes were broadly similar for both samples. Figs. 7b,c provide a side-by-side comparison of the MICrONS patterns obtained using the unweighted definition of the synaptic pattern (from Figs. 2–4) and the weighted definition based on Eq. (1). The weighted formulation is intended to compensate for potential non-uniformities in sample geometry. Considering the top view of the MICrONS cortical column (Fig.7a) and the corresponding two-dimensional projections, only minor differences are observed between the two approaches—except perhaps in Layer 5, where the discrepancy is somewhat more noticeable, though the overall pattern shape remains consistent. Both methods thus yield comparable results, indicating that the spatial sampling density does not vary systematically with direction; in other words, sampling is sufficiently isotropic that a computationally intensive weighting correction is not required.

**Figure 7.**
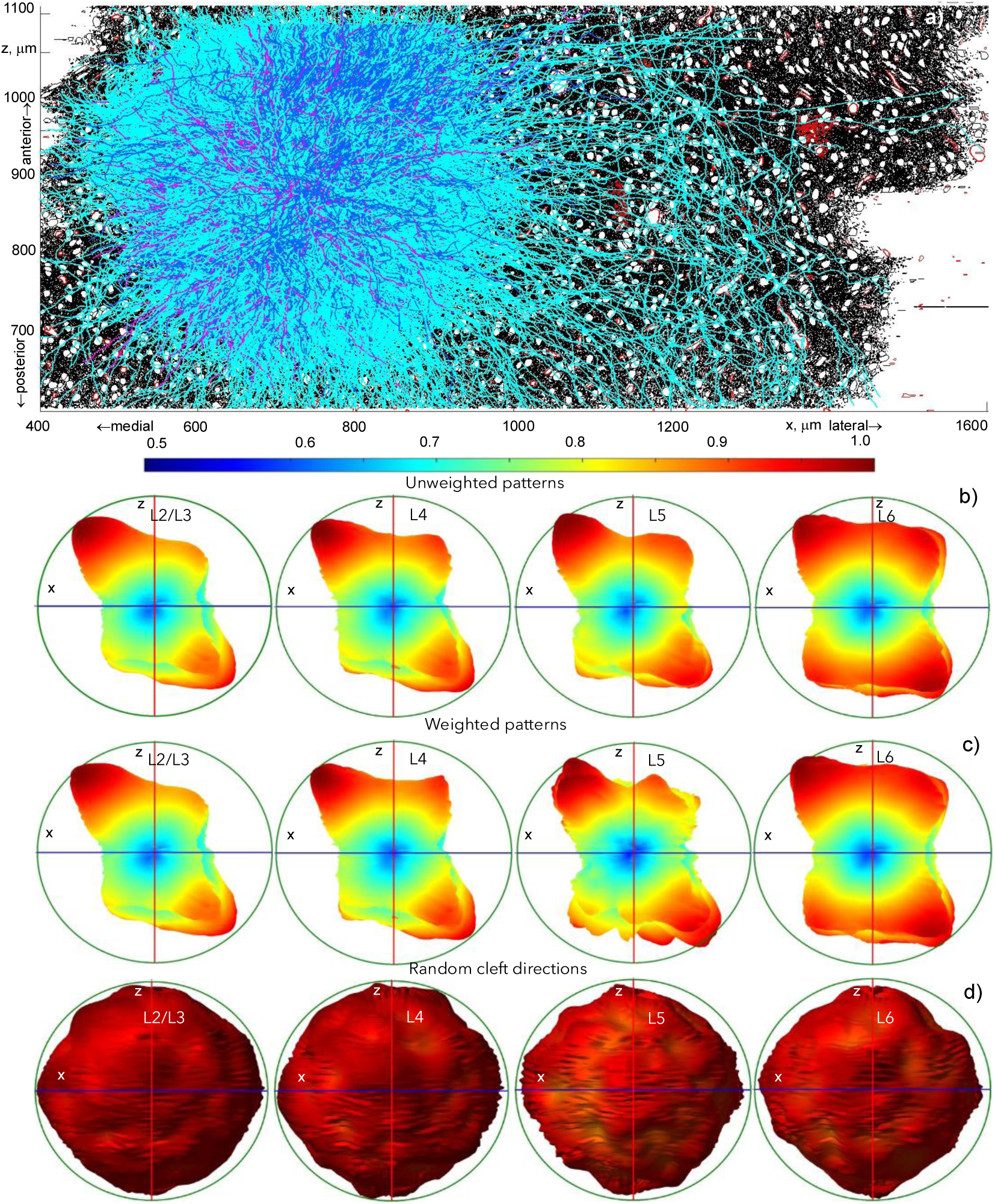
a) Top view (into layers) of IARPA Phase III MICrONS V1 cortical column with 1,097 excitatory neurons. Axons – cyan, apical dendrites – magenta, basal dendrites – blue. The average length of the axonal arbor per neuron is 4.9 mm. The column overlaps with the original Minnie 65 sample’s cross-section at y=600 μm where cell membranes are shown in black, the blood network – in red, and the somas are white. b). Top view (xz -plane into the cortical column) of the same sample. c, d) Side-by side comparison between the patterns for the MICrONS sample obtained with the unweighted definition of the synaptic pattern (from Figs. 2,3,4) and using the weighted definition from Eq. (1), respectively. The same xz-view is displayed. e) The same xz-view of synaptic patterns obtained when all synaptic cleft directions are assigned random values chosen from the 3D standard normal distribution with mean zero and standard deviation one.

### 4.5 Randomized controls

As an additional control, we assigned random orientations to all synaptic clefts, drawing each direction from a three-dimensional standard normal distribution (mean = 0, standard deviation = 1). This procedure was applied to both the mesh-touch human synaptic patterns and the mouse synaptic patterns. The resulting distribution for the mouse cortical column is shown in Fig.7d. In every such randomized dataset, the resulting patterns were nearly omnidirectional, as illustrated in Fig.7d. The corresponding human results were similar.

### 4.6 Known (major) artifact(s)

Fig.8 explains why the apparent strong minimum in the anterior–posterior (*z*) direction observed in the voxel-based patterns of the H01 sample (Figs. 2d, 3d, and 4d of the main text) arises from methodological artifacts rather than a genuine biological feature. Fig.8a shows three consecutive H01 slices containing a synapse oriented primarily within the *xy*-plane (highlighted by yellow circles). This synapse is readily detected during manual labeling and classifier training on electron microscopy cross-sections, which have high in-plane (*x* − *y*) resolution of 4 nm but lower axial (*z*) resolution of 33 nm. In contrast, Fig.8b shows three consecutive H01 slices containing a synapse oriented predominantly along the *z*-axis. Such a synapse is often visible in only a single *z*-slice, resulting in poor detection and contributing to their apparent underrepresentation.

**Figure 8.**
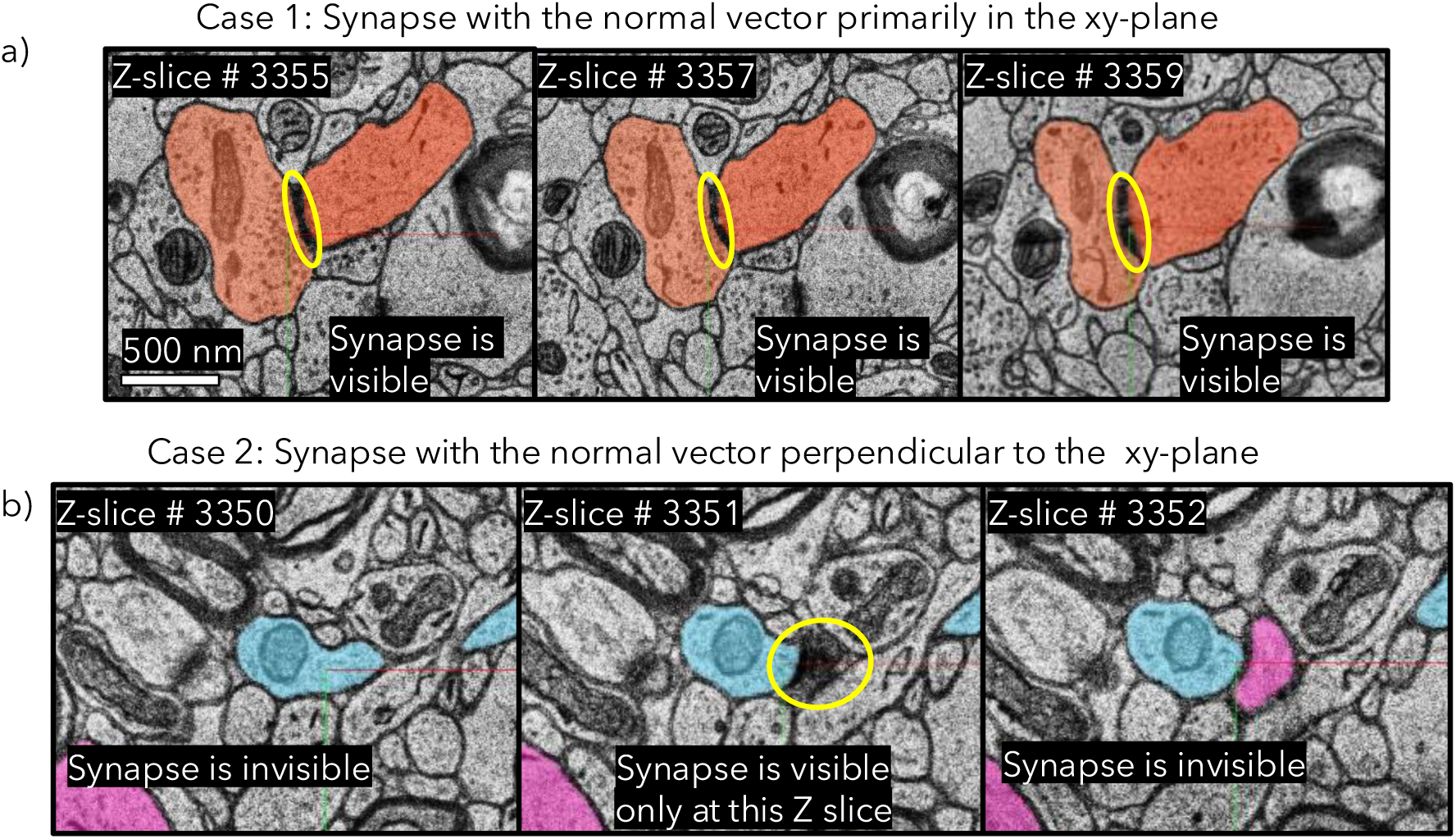
H01 synapses oriented primarily within the *xy*-plane (highlighted by yellow circles) are readily detected during manual labeling and classifier training on electron microscopy cross-sections, which have high in-plane (*x* − *y*) resolution of 4 nm but lower axial (*z*) resolution of 33 nm. (b) In contrast, H01 synapses oriented predominantly along the *z*-axis are often visible in only a single *z*-slice, resulting in poor detection and contributing to their apparent underrepresentation.

A comparable artifact may also occur in the MICRONS cortical column, which was imaged at a similarly anisotropic resolution. The synaptic patterns in Figs.2g,3g,4g of the main text and in Fig.7b,c display corresponding minima that align closely with the Cartesian axes.

## Acknowledgments

SNM has received support from NIH grant R01MH130490, R01EB035484; BD has received support from NIH grant U24NS120053; ZDD has received support from NIMH grant ZIAMH002955; HL has received support NIDA grant ZIADA000638; MB has received support from NIDA grant UG3DA048502, NIGMS grant T34GM137858, T32GM136499, NINDS grant R01NS112996, R01NS101362; DB has received support from NIMH grant UG3MH123386, NINDS grant U19NS104653, U24NS109102, NIBIB grant U01EB026996; BR has received support from NIBIB grant P41EB030006.

## Notes

### Competing Interest Statement

The authors have declared no competing interest.

### Summary of Updates

Updated Discussion section to include sub-headings and some additional explanation of possible mechanistic basis. References were updated with additional items

